# Native *Olympia* oysters in San Francisco Bay: microbiome and antimicrobial resistance derived insights into restoration

**DOI:** 10.1101/2025.10.30.685688

**Authors:** Nishta Mukundan, Sanjiev Nand, Archana Anand

**Author notes:** Equal contributing authors (Nishta Mukundan, Sanjiev Nand).

## Abstract

Restoration of Olympia oyster (*Ostrea lurida*) reefs can deliver ecological and public-health co-benefits. Characterization of the microbiome can provide valuable insights towards reef health but remains underused especially in San Francisco Bay. To overcome this gap, we applied shotgun metagenomics to oyster tissue, water, sediment, and biofilm collected from five San Francisco Bay sites (Loch Lomond, Point San Quentin, Brickyard Park, China Camp, Dunphy Park; Nov 2023) to profile community structure, Vibrio diversity, and antimicrobial-resistance (AMR) potential. Evaluating beta-diversity based on Bray–Curtis dissimilarities, we saw strong clustering by sample type (PERMANOVA R^2^ = 0.645, p = 0.001) and no significant clustering by location (R^2^ = 0.068, p = 0.937). Alpha diversity (Shannon) differed by matrix (Kruskal–Wallis p = 0.000416), with sediment and water highest and oyster tissues lowest. Vibrio taxa occurred at low relative abundance across matrices but were slightly enriched in biofilms and tissues; several Vibrio phages were also detected, consistent with potential (but unconfirmed) top-down control. Resistome profiling revealed a long-tailed distribution dominated by the tetracycline efflux pump tet(C) (70,049 reads) and APH(3⍰)-Ia (4,498 reads), alongside multidrug efflux components (acrAB/-D, mdtABC/H/F/P, mdfA, MexF/MexW). Biofilms were AMR hotspots, while oyster tissues uniquely harbored Vibrio cholerae varG (Subclass B1 β-lactamase). Collectively, our results indicate that matrix, not geography, structures reef microbiomes, that *Vibrio* is present but scarce, and that biofilms concentrate AMR potential. We propose a microbiome-integrated monitoring approach that stratifies by matrix, prioritizes biofilm resistome tracking, and targets Vibrio surveillance in oysters to align restoration performance with ecosystem-health safeguards.

## 1. INTRODUCTION

Historically, oysters have played a critical role in stabilizing ecosystems and preserving environments (Zu Ermgassen et al., 2012). Oysters fill the role of keystone species, and as such, their population dynamics directly influence the relative abundance and prosperity of various associated marine organisms (Rajagopal et al., 2008). In addition, oyster reefs provide natural shoreline protection by reducing wave strength, both mitigating erosion and dampening the impact of storm surges (Boyer et al., 2017; Gedan et al., 2010; Chowdhury et al., 2023). Highly complex and durable reef structures also provide a crucial buffer against extreme weather events that are increasing in frequency due to climate change. Beyond regulatory services, reefs exist as complex habitats that support a diverse range of marine species (Coen et al., 2007; Chan et al., 2022), contributing to global biogeochemical cycles by facilitating organic carbon storage and cycling, specifically through the sequestration of blue carbon (Fodrie et al., 2017). Oysters are also significant contributors to water quality through their crucial role in water filtration. A single adult oyster can filter over 50 gallons of water per day, removing excess particulates and elements that may be harmful to coastal systems (NOAA, 2020; Kellogg et al., 2014). Through such filtration and nutrient sequestration, oyster populations can not only reduce cultural eutrophication, but also serve to improve ecosystem function as a whole (Bricker et al., 2019).

The Olympia oyster (*Ostrea lurida*) is the only oyster species native to the U.S. West Coast. Once abundant, its populations have declined significantly since the mid-19th century Gold Rush due to centuries of overfishing, habitat destruction, sedimentation, and nutrient loading (Kirby 2004). Prior to this drastic population decline, Olympia oysters were consumed in substantial quantities by Native Americans and settlers (Ridlon et al., 2021). Excess nutrient inputs have caused eutrophication of resident water bodies, resulting in frequent algal blooms and hypoxic zones that further stress remnant populations (Wasson et al., 2014). Today, most existing Olympia oyster beds in the San Francisco Bay are restored rather than natural, as historic reefs were nearly wiped out during early exploration. Given the species’ ecological services of shoreline stabilization, habitat creation, and water filtration, restoration of Olympia oyster beds is a priority for improving ecosystem function and health in the Bay. Studies have been conducted highlighting the positive impacts of restoration, showing that such beds can provide equal ecological benefits as natural ones (Smith, Cheng, & Castronati 2022). As such, the restoration of oyster beds in the San Francisco Bay is critical to preserving the well-being of coastal ecosystems.

Previous restoration initiatives in the Bay, such as the San Francisco Bay Living Shorelines Project and the Native Olympia Oyster Collaborative, have primarily focused on physical and ecological metrics including reef structure, substrate stabilization, and population density (Ridlon et al., 2021). Although these metrics are robust, restoration efforts continue to struggle with sustaining population densities. Emerging evidence shows that oysters host complex microbiomes that shape host fitness, ecosystem function, and water quality (Lokmer & Wegner, 2015), yet microbial processes remain comparatively understudied in restoration design. Analogous work in coral reef restoration demonstrates that host-associated microbiomes are integral to reef health and can serve as indicators of environmental change. A recent study (Cyphert et al., 2024) addresses this gap by characterizing oyster-associated and oyster bed-adjacent microbial communities across Bay sites and conditions, identifying candidate microbial indicators of reef performance and pathogen risk, and recommending a microbiome-integrated framework to guide future restoration and monitoring.

Building on these microbiome indicators, we next consider antimicrobial resistance (AMR) as a parallel axis of reef performance and human-health risk. Antimicrobial resistance (AMR) in marine environments is rising globally (Kumarage et al., 2022; Dutta et al., 2021). Against this backdrop, oyster-associated and reef-adjacent microbiomes may function as reservoirs of AMR genes with implications for restoration outcomes and human health. Of particular concern are interactions with *Vibrio*, a genus common in coastal waters containing pathogenic species such as *V. parahaemolyticus* and *V. vulnificus*, which cause seafood-borne gastroenteritis and, in severe cases, septicemia (Froelich & Noble, 2016). Because oysters filter large volumes of water, they can concentrate *Vibrio* and potentially act as vectors of human exposure (Diner et al., 2023). At the same time, commensal microbes associated with oyster tissues can suppress *Vibrio* via competitive exclusion and other antagonistic mechanisms (Lokmer et al., 2016). While these dynamics have been explored in Pacific oysters (*Crassostrea gigas*; Wang et al., 2021), they remain largely uncharacterized in the native Olympia oyster (*Ostrea lurida*) as overfishing and legacy mercury contamination has made the harvesting and consumption of Olympia oysters largely obsolete in the San Francisco Bay (Kolipinski et al., 2020). Clarifying these interactions is critical for designing microbiome-aware restoration strategies that bolster oyster and reef health while reducing Vibrio-related risk. To prevent future exposure and the concatenation of harmful pathogens, it is important to consider competitive exclusion mechanisms and exposure potential when informing restoration efforts. Restoration informed on microbial dynamics will create more sustainable, healthy reefs with the potential for future harvesting and ecological benefits.

To address this knowledge gap, we conducted a shotgun metagenomic profiling of Olympia oyster tissues and nearby water, sediment, and biofilm communities across San Francisco Bay. We quantified community composition with an emphasis on *Vibrio* diversity, and we screened for AMR genes to evaluate potential reservoirs and co-occurrence patterns relevant to pathogen control. Our findings demonstrate how microbiome characterization can inform adaptive restoration and concurrently mitigate human-health risks linked to bacterial pathogens.

## 2. METHODS

### 2.1. Site Description and Sample Processing

Data was collected in November 2023 from five San Francisco Bay sites (Fig.1): Loch Lomond, Point San Quentin, Brickyard Park, China Camp, and Dunphy Park. Dunphy Park, Brickyard Park, and the area near Point San Quentin are established *Ostrea lurida* restoration sites, while China Camp hosts a natural *Ostrea lurida* population that survived early exploration. Loch Lomond was cited as a high-potential candidate for conservation with high adult oyster density metrics (Wasson et al., 2014). At each site, sediment was sampled into three 50mL Falcon tubes from the upper 10cm and placed in sterile bags. Biofilms were scraped into three 15mL Falcon tubes prefilled with 99.5% ethanol. Oysters (≥5 individuals per site) were collected and sealed in separate sterilized bags. All samples were kept on ice in a cooler for transport to San Francisco State University and stored at -80°C until downstream processing (Cyphert et al., 2024).

**Figure 1.**
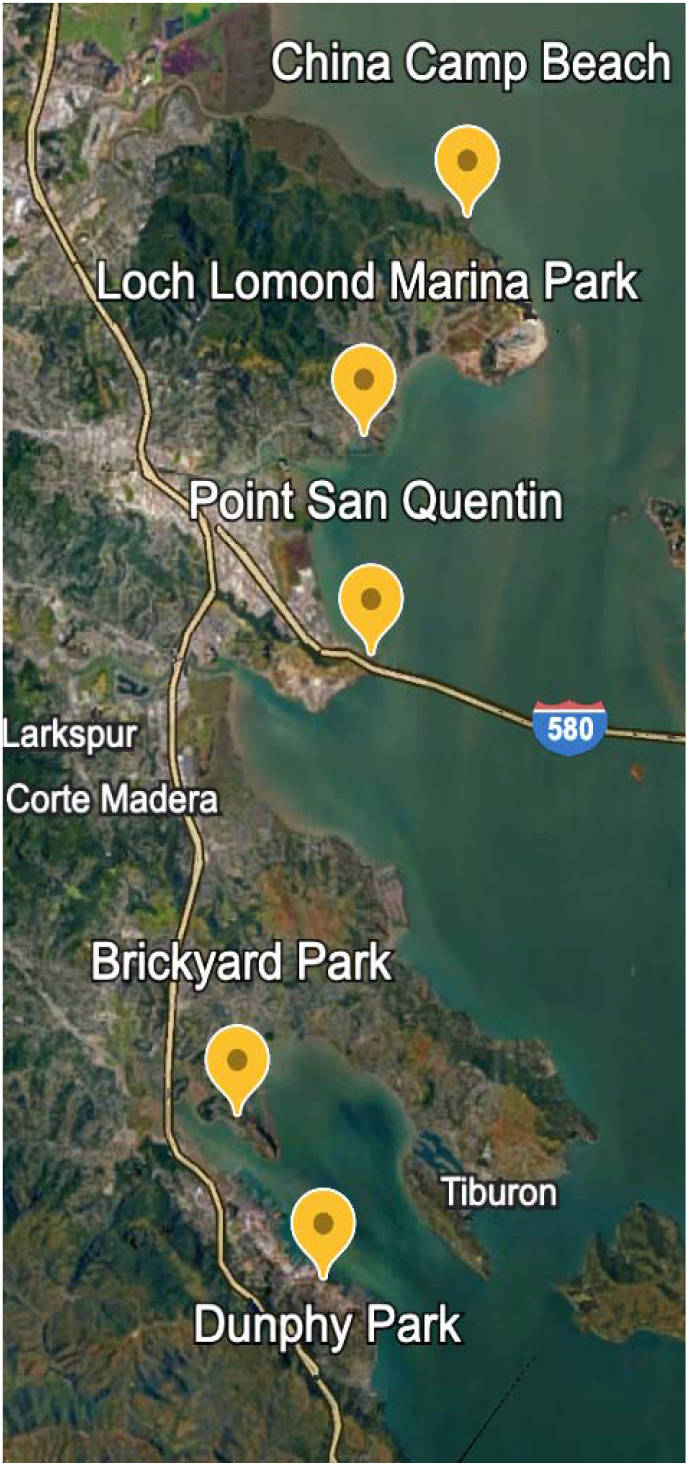
Restored oyster beds across the San Francisco Bay Area where oyster tissue, sediment, water, and biofilm were sampled

### 2.2. Nucleic Acid Extractions and Processing

DNA extraction followed the methods detailed in Cyphert et al., 2024 wherein oyster tissue DNA extraction saw 25 mg ± 1 mg of oyster tissue incubated in a water bath at 56°C overnight (12+ hrs.), and soil, and biofilm DNA extractions were conducted using the DNeasy PowerSoil Pro Kit according to the manufacturer’s instructions. Extracted DNA was stored at −80°C until sequencing. Paired-end illumina shotgun sequencing (30-50M read depth) was performed on the samples to generate fastq files that could be interpreted using statistical methods.

### 2.3. Bioinformatics and Statistical Analysis

Sequence data was analyzed using the CZID (previously IDseq) platform. Raw sequencing data was processed using CZID’s illumina pipeline, an open-source, cloud-based pipeline for metagenomic analysis (Kalantar et al., 2020). CZID performs host filtering, quality control, alignment, and taxonomic classification services, returning microbial abundance tables for each sample. AMR data was also processed through the CZID pipeline (Alcock et al., 2019). Statistical analyses were performed using RStudio (RStudio 2021). Alpha diversity calculations (Shannon index, Richness) were performed using a one-way analysis of variance (ANOVA) that yielded significance across sample type and location, while Bray-Curtis beta diversity was assessed using a permutational multivariate analysis of variance (PERMANOVA; adonis2 function) which calculated variance in microbiota composition by sample type and location (Fig2) (Anderson 2017; Cyphert et al., 2024).

**Figure 2.**
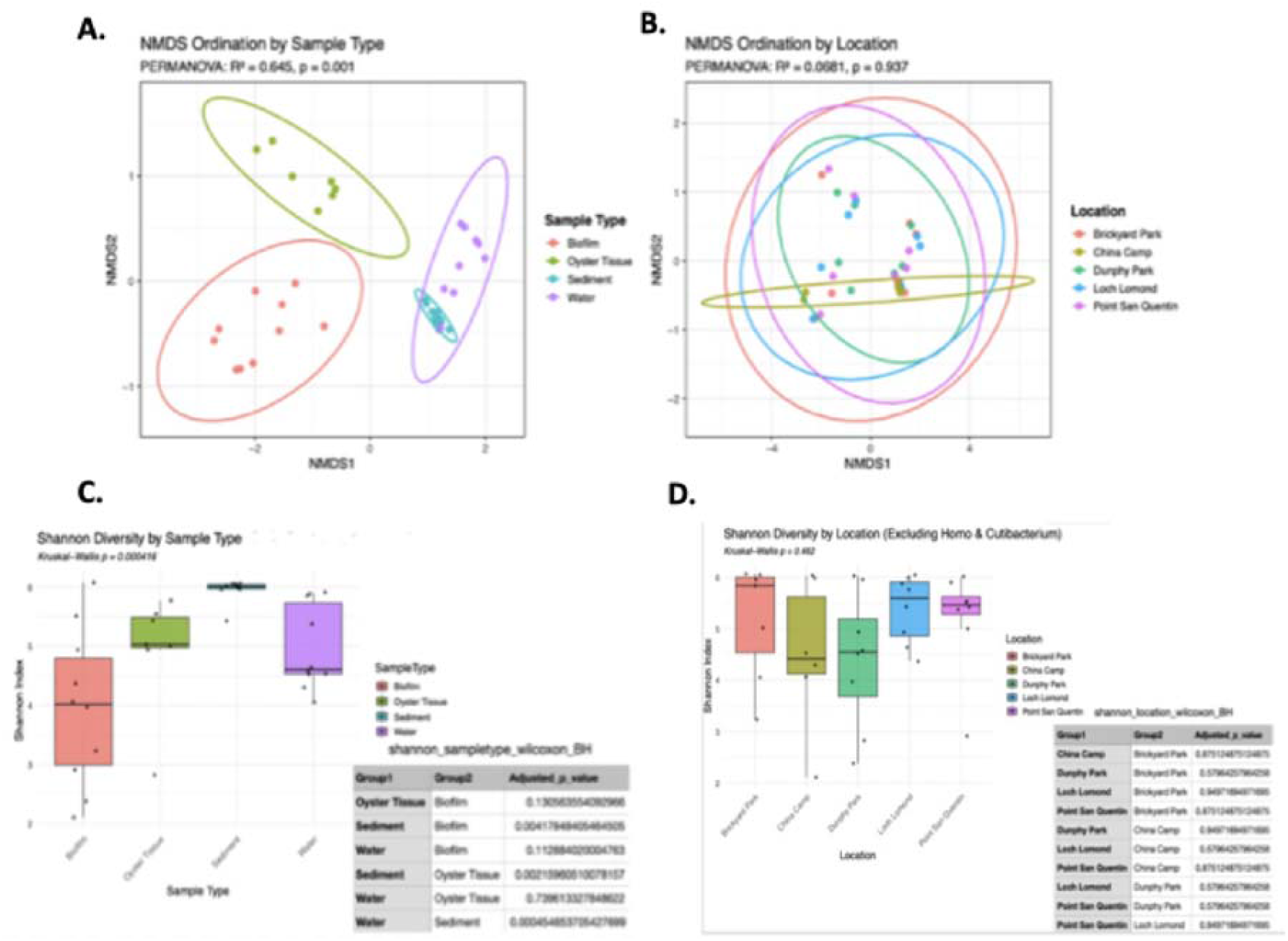
General Structure, Diversity, and Alpha Diversity of microbial communities by Sample Type and Location. NMDS ordination plots based on Bray-Curtis dissimilarity of genus-level microbial communities by sample type show strong sample clustering (PERMANOVA R 2 = 0.645, p= 0.001) (A), whereas NMDS ordination by location shows no significant clustering (p=0.937) (B). Shannon diversity differed significantly by sample type (Kruskal-Wallis p=0.000416), with sediment and water samples showing the highest diversity and oyster tissue the lowest (C). Shannon diversity by location showed overall group differences (Kruskal– Wallis p= 0.042), but no significant pairwise contrasts (D).

CZID raw NTR data and associated sample metadata was organized into the following categories: Top 20 Taxa by site (Fig4A), and by sample type (Fig4B) using R. Similarly, top 20 Vibrio and known AMR sequence data was also sorted by site (Fig4C) and sample type (Fig4D) using R software. Finally, these curated data sets were used to pull out and sort Vibrio associated AMR sequences from the oyster microbiomes using R statistical analysis as well (Fig5).

**Figure 4.**
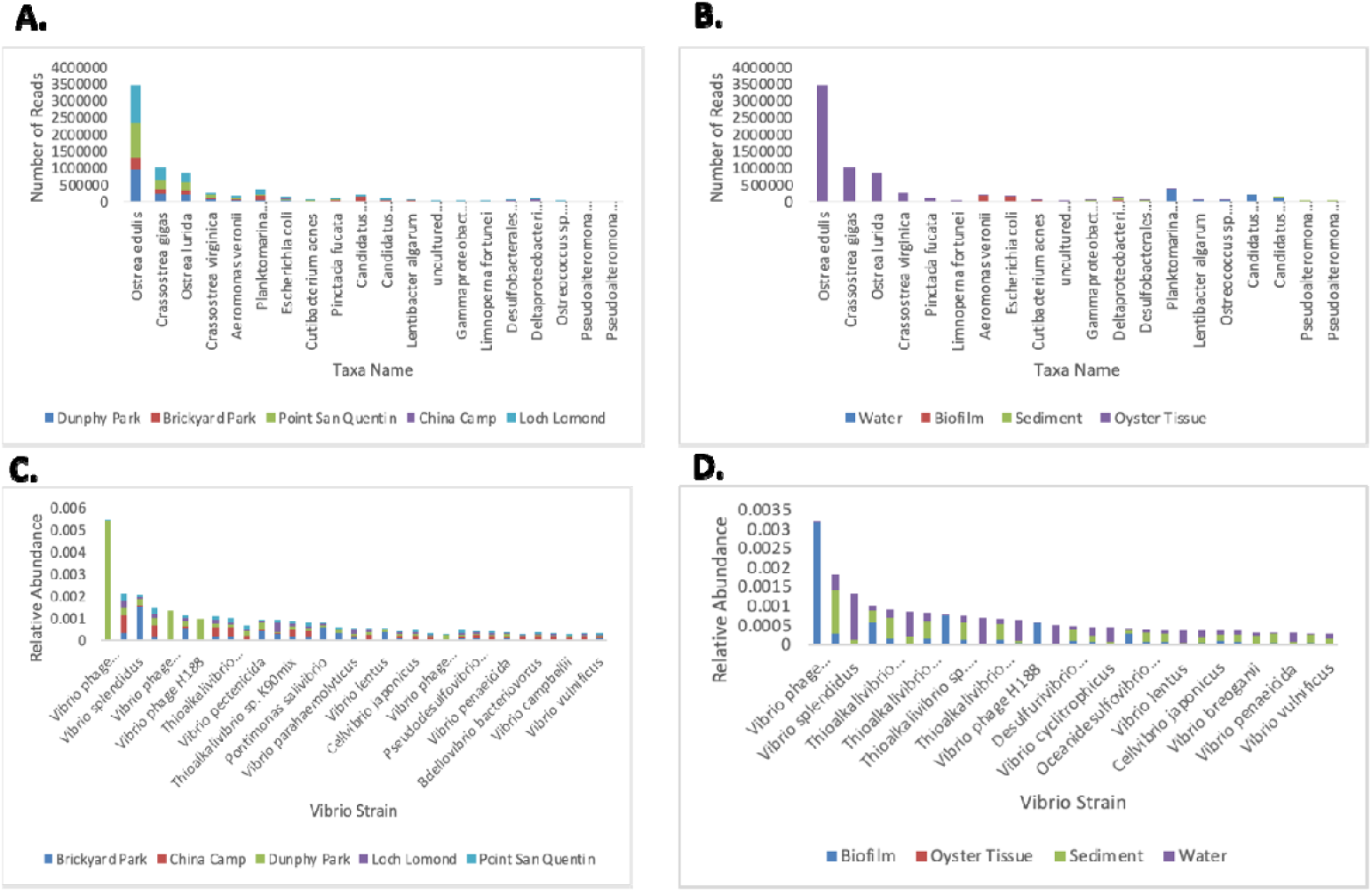
Microbial Abundance in Oyster beds measured by observing abundance of top 20 taxa by location (A), and by sample type (B) as well as relative abundance of top 20 *Vibrio* taxa by location (C), and sample type (D). Reveals high taxonomic prevalence of oyster tissues (especially *Ostrea edulis*), as well as limited but still existent presence of *Vibrio* and associated *Vibrio* phages, potentially indicative of pathogen control mechanisms through competition.

**Figure 6.**
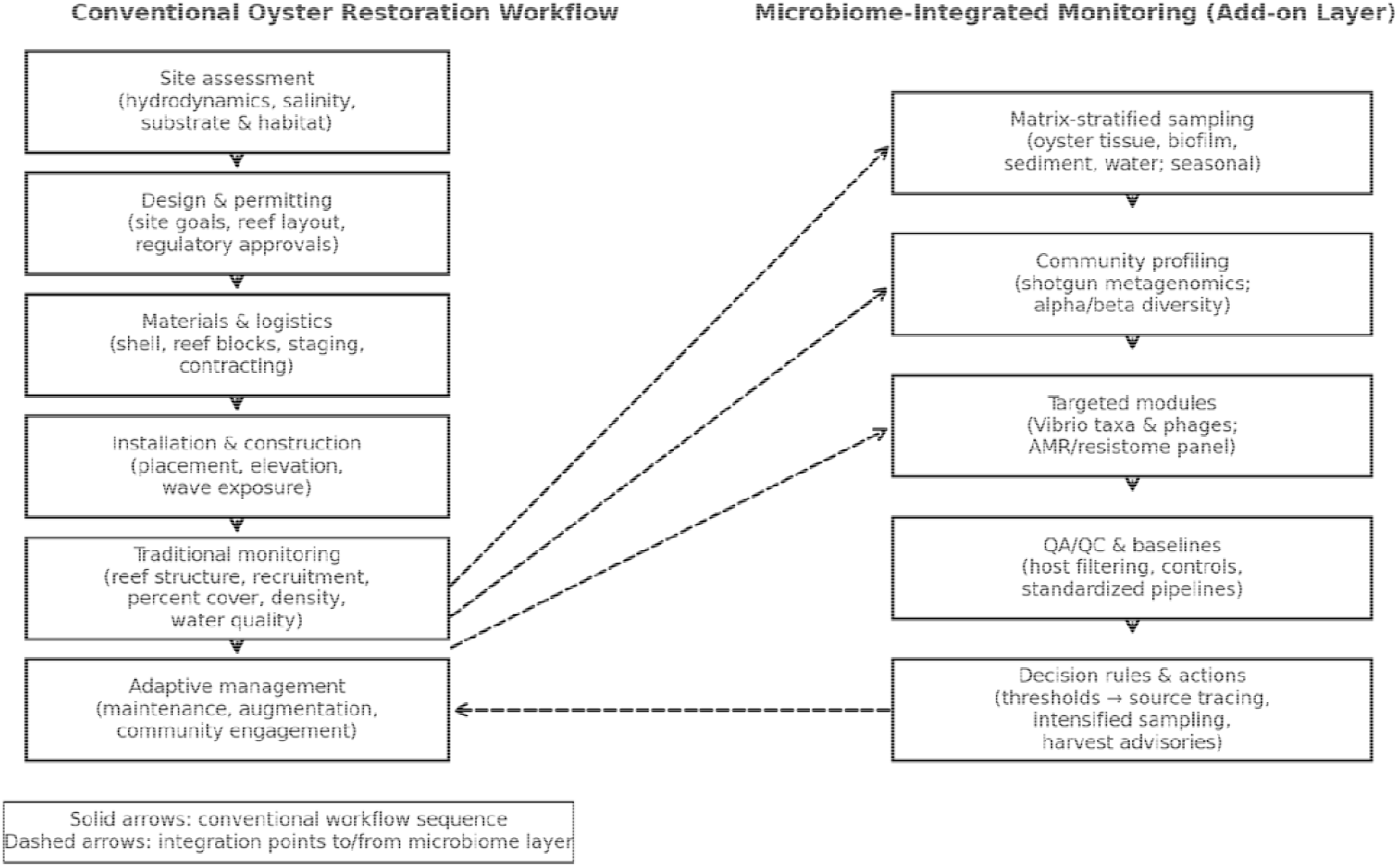
Integrated framework for oyster reef restoration. Left: conventional workflow (site assessment → design & permitting → materials/logistics → installation → traditional monitoring → adaptive management). Right: microbiome-integrated monitoring layer added at the monitoring stage, including matrix-stratified sampling (oyster tissue, biofilm, sediment, water), community profiling, targeted Vibrio/AMR modules, QA/QC and baselines, and decision rules that feed back into adaptive management. Solid arrows show the standard sequence; dashed arrows show integration points and feedback loops.

## 3. RESULTS

To quantify ecological and health-relevant signals, we profiled community composition, *Vibrio* prevalence, and AMR gene carriage across sites and sample types. Microbial community structure was driven by sample type rather than site. NMDS ordination based on Bray–Curtis dissimilarities showed strong clustering by sample type (PERMANOVA R^2^ = 0.645, p = 0.001; Fig. 2A), whereas samples did not cluster by location (R^2^ = 0.068, p = 0.937; Fig. 2B). Alpha diversity (Shannon) differed significantly among sample types (Kruskal–Wallis p = 0.000416), with sediment and water exhibiting the highest diversity and oyster tissue the lowest (Fig. 2C). Diversity varied across locations overall (Kruskal–Wallis p = 0.042) but without significant pairwise contrasts (Fig. 2D). Taxonomic profiles were dominated by host-associated reads and common estuarine taxa; *Vibrio* reads were consistently low in relative abundance across sites and matrices, with slightly higher values in biofilms and oyster tissues than in sediments and water (Fig. 4C–D). AMR screening recovered a long-tailed distribution of resistance determinants, with the tetracycline efflux pump tet(C) overwhelmingly predominant (70,049 reads) and APH(3⍰)-Ia also common (4,498 reads), followed by lower-abundance multidrug efflux components (e.g., mdtABC, acrAB, AcrF) (Fig. 3). Matrix-resolved hits indicated biofilms as hotspots for tet(C) and APH(3⍰)-Ia, whereas oyster tissues uniquely contained Vibrio cholerae varG (Subclass B1 β-lactamase), suggesting potential carriage of *Vibrio*-linked resistance in the host compartment (Fig. 5).

Collectively, these results indicate that (i) sample type is the principal axis of microbiome variation in restored Olympia oyster reefs, (ii) *Vibrio* taxa are present at low relative abundance, and (iii) AMR genes are most enriched in biofilms, and we also detect some Vibrio-linked resistance markers in oyster tissues.

## 4. DISCUSSION

This study leveraged shotgun metagenomics to evaluate microbiome structure, *Vibrio* dynamics, and antimicrobial-resistance (AMR) potential across restored Olympia oyster (*Ostrea lurida*) reefs in San Francisco Bay. Beta-diversity analyses showed that community composition was structured primarily by sample type (oyster tissue, water, sediment, biofilm) rather than site, with strong clustering by matrix (PERMANOVA R^2^ = 0.645, p = 0.001) and no significant clustering by location. Alpha diversity likewise differed by matrix, with higher Shannon diversity in sediment and water and lower diversity in oyster tissues. These patterns echo prior works, where host association and habitat often outweigh geography in explaining microbiome variation (Stevick et al., 2021; Diner et al., 2023) and suggest that restoration monitoring can generalize across the Bay Area when stratified by matrix..

Host-associated reads were abundant in tissue libraries, including assignments to Ostrea spp. Notably, we observed a high presence of *Ostrea edulis* when analyzing oyster tissue samples (Fig4). *O. edulis* have been proven to be highly beneficial in ecosystem stabilization (Rodriguez-Perez et al., 2019; Kennon et al., 2023), indicating that they may have been imported and used in early restoration efforts as a means of stabilizing restored communities. However, we note that high host read proportions on this type of oyster (*O*.*edulis* genomes) can inflate apparent host signal in metagenomic classifications such as CZID, thus, these assignments likely reflect host enrichment rather than non-native species presence (Bean et al., 2022; Siegwald et al., 2017). Regardless, restored sites exhibited mixed consortia of host-associated, sediment-associated, and free-living taxa, consistent with multi-habitat coupling expected on mature reefs (Johnson et al., 2024). Further exploration into the reasons behind observation of *O. edulis* within *O. lurida* samples would shed further light on this phenomenon.

Targeted inspection of *Vibrio* strains indicated low relative abundance across sites and matrices with slightly higher representation in biofilms and tissues. The dominant taxa across restoration sites were Vibrio spp. (including *V. crassostrea, V. splendidus*, and *V. parahaemolyticus*) and Thioalkalivibrio species. Several *Vibrio* phages (ex. Vibrio phage 1.087.A._10N.261.45.F9) were among the top *Vibrio*-associated taxa, accompanied by phage counterparts targeting Vibrio (Fig4). This community composition aligns with microbial profiles observed in sediments and biofilms sampled in existing papers that focus on *Vibrio* (Brossard Stoos et al., 2022), reflecting an active nutrient cycling network typical of mature restoration environments. The presence of *Vibrio* phages is indicative of potential top-down control mechanisms that may counter pathogenic Vibrio populations posing threats to human health (Bruto et al., 2016). Most of these phages were observed in biofilm samples (Fig4D), indicating a healthy bacterial community. However, this observation may potentially indicate environmental stress that favors *Vibrio* growth (Brumfield et al., 2023), or host-pathogen co-evolution facilitated by phage presence (Piel et al., 2022). BLAST analysis of the recorded phages was inconclusive regarding lifestyle, returning no clear integrase or lysis annotations. However, without definitive lifestyle annotations or cultivation data, these should be viewed as candidate indicators rather than proven antagonists. Future studies that aim to conduct plaque morphology tests would be helpful experiments to determine the lifecycle (lytic vs. temperate) of the observed phages before making conclusive decisions about their role in *Vibrio* control or propagation.

Analysis of the top 20 antimicrobial resistance (AMR) genes revealed a significant presence of multidrug efflux pumps and stress response genes, including acrB, acrD, mdtA, mdtP, mdtM, MexF, MexW, and mdfA (Table 1). Efflux pumps export antibiotics and other metabolites contributing to broad resistance phenotypes (Blanco et al., 2016) and promoting biofilm formation, which can reinforce community persistence (Alav et al., 2018). Their prevalence suggests potential competitive exclusion mechanisms that may suppress *Vibrio* colonization by outcompeting *Vibrio* stains for resources in biofilm and sediment microenvironments (Eickhoff & Bassler, 2021). Their prevalence especially in biofilms supports a role for competitive exclusion that may limit *Vibrio* colonization by outcompeting *Vibrio* strains for space and resources in biofilm and sediment microhabitats. Consistent with coral reef studies in which microbial composition tracks reef health, environmental stress, resilience, and pathogenic prevalence (Li et al., 2022),these microbial structures and AMR related genes could be useful for diagnosing environmental stress, improving the quality of oyster restoration efforts.

**Table 1.** AMR genes dominated by tetracycline efflux pump tet(c) (70049 reads) as well as APH(3’)-la (4498 reads). Also shows presence of various multidrug efflux pumps indicating potential control mechanisms and mature bacterial communities but perhaps has other indicators for reef restoration and resistance control.

| Gene Name | # of Reads |
| --- | --- |
| tet(c) | 70049 |
| APH(3')-Ia | 4498 |
| Bifidobacterium adolescentis rpoB mutants conferring resistance to rifampicin | 628 |
| rpoB2 | 552 |
| mdtB | 244 |
| mdtC | 200 |
| acrD | 158 |
| acrB | 128 |
| mdtA | 98 |
| AcrF | 82 |
| Escherichia coli mdfA | 80 |
| mdtH | 70 |
| bacA | 58 |
| MexW | 55 |
| Bifidobacterium bifidum ileS conferring resistance to mupirocin | 55 |
| kdpE | 52 |
| mdtF | 50 |
| mdtP | 50 |
| mdtM | 50 |
| MexF | 49 |

Detection of bacA, a bacitracin resistance conferring gene, indicates the possible presence of Bacillus or related taxa that are capable of producing peptide antibiotics (Sermkaew et al., 2024). These antibiotics are active against Gram-negative bacteria (such as *Vibrio*) that are typically more resistant to antibiotics and varying antimicrobial agents compared to their positive counterparts (Barreto-Santamaría et al., 2021). The possibility of antibiotic generation has various implications for human health, perhaps providing mechanisms for countering virulent *Vibrio* and other pathogenic strains that pose dangers to humans via consumption.

Further, detection of tet(C), a tetracycline resistance gene, and APH(3⍰)-Ia (Table 2), an aminoglycoside resistance gene, point to the possibility of horizontal gene exchange in the environment, suggesting that resistance traits may thrive under man-made selection pressures that are far more common in restoration settings (Tao et al., 2022). These genes have the potential to enhance overall microbiota resilience, perhaps limiting Vibrio colonization indirectly through competition, reflecting recent research detailing how efflux-facilitated resistance contributes to the overall ecological wellbeing of environmental microbiomes, and can shape microbial dynamics in aquatic systems (Stephen & Thomas, 2022). However, overaccumulation of these genes may provide opportunistic pathogens with various resistance mechanisms through horizontal gene transfer, posing threats to human health. This possibility further stresses the importance of regulating microbial environments in restoration sites to prevent dangerous microbial imbalances. Tet(c) prevalence also suggests persisting antibiotic influence in the surrounding environment, consistent with prior studies describing tet(C) enrichment in coastal sediments near restoration zones (Roberts & Schwarz, 2016). Future work should aim to couple microbial analysis with co-occurrence network approaches to truly determine whether efflux related AMR genes are actively contributing to *Vibrio* suppression.

**Table 2.**
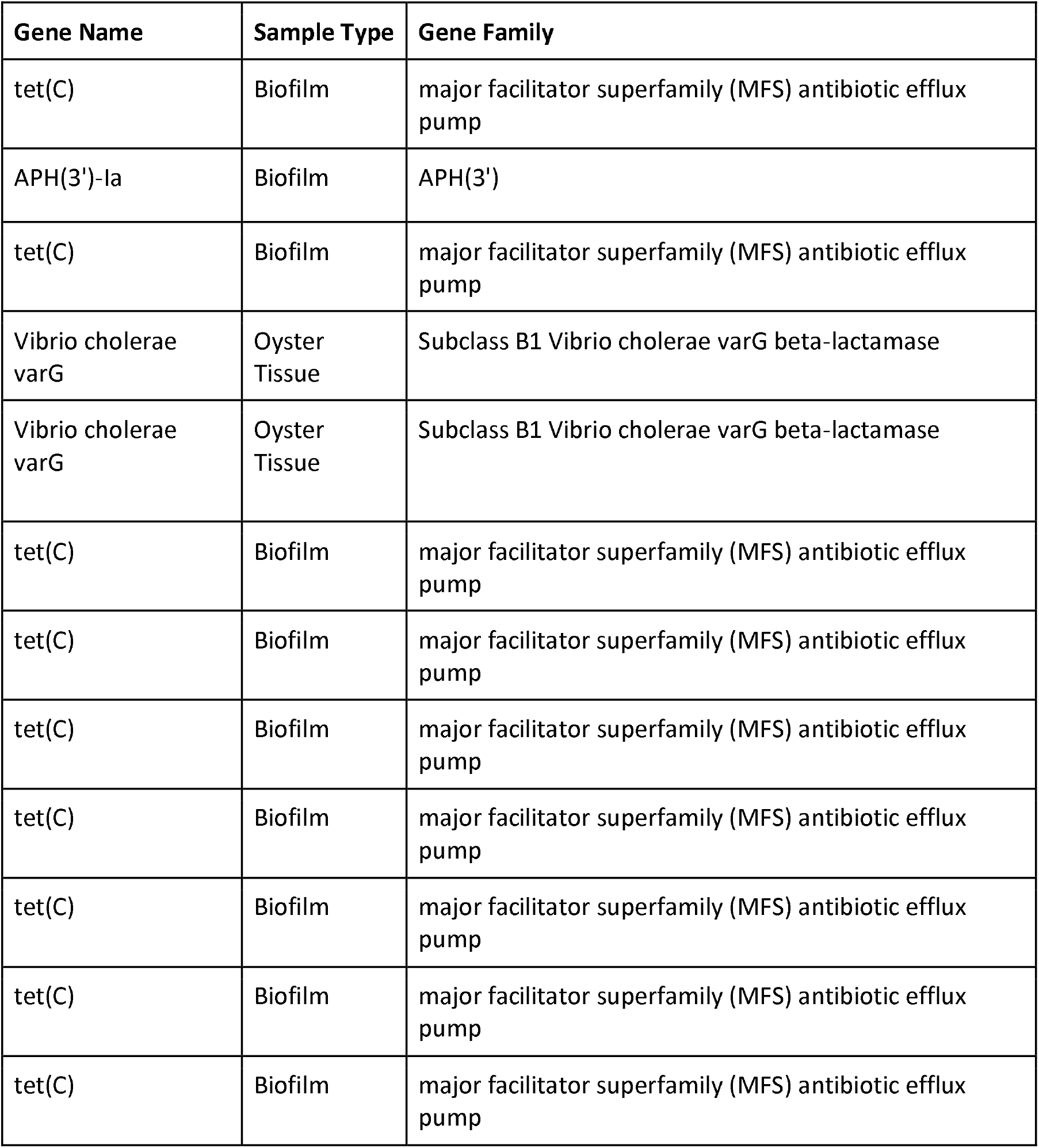
Vibrio related AMR genes found in samples reveal a high prevalence of tet(C), an antibiotic efflux pump with potential for competitive exclusion. Also reveals high prevalence of AMR genes in biofilms as opposed to other sample types. Noted is the presence of Vibrio cholerae varG in oyster tissue samples, potentially hinting towards resistant or highly durable *Vibrio* strains that may need to be monitored in restoration sites to avoid concerns for human health.

| Gene Name | Sample Type | Gene Family |
| --- | --- | --- |
| tet(C) | Biofilm | major facilitator superfamily (MFS) antibiotic efflux pump |
| APH(3')-Ia | Biofilm | APH(3') |
| tet(C) | Biofilm | major facilitator superfamily (MFS) antibiotic efflux pump |
| Vibrio cholerae varG | Oyster Tissue | Subclass B1 Vibrio cholerae varG beta-lactamase |
| Vibrio cholerae varG | Oyster Tissue | Subclass B1 Vibrio cholerae varG beta-lactamase |
| tet(C) | Biofilm | major facilitator superfamily (MFS) antibiotic efflux pump |
| tet(C) | Biofilm | major facilitator superfamily (MFS) antibiotic efflux pump |
| tet(C) | Biofilm | major facilitator superfamily (MFS) antibiotic efflux pump |
| tet(C) | Biofilm | major facilitator superfamily (MFS) antibiotic efflux pump |
| tet(C) | Biofilm | major facilitator superfamily (MFS) antibiotic efflux pump |
| tet(C) | Biofilm | major facilitator superfamily (MFS) antibiotic efflux pump |
| tet(C) | Biofilm | major facilitator superfamily (MFS) antibiotic efflux pump |

Additionally, it can be inferred that the microbial environments observed in the sampling are active and adaptive due to the detection of regulatory elements such as kdpE (Table 1), which increases potassium uptake in high stress situations (Freeman et al., 2013), and rpoB variants (Table 1) associated with rifampicin resistance which could be an indicator for multidrug resistant *Staphylococcus* spp. among other pathogens (Edslev et al., 2025). Consideration of these microbes is critical for adequate restoration, as observing kdpE concentration and behavior can be indicative of environmental stress and overall reef health. Observation of such regulatory elements can help inform restoration initiatives on the needs of reefs, helping to address immediate stressors and improve restoration quality. These AMR profiles depict a resilient community that has the potential to not only weather environmental disruptions, but also inhibit opportunistic pathogen colonization through competitive exclusion, biofilm formation, and the production of antimicrobial compounds. Monitoring these diverse microbial communities could provide signals for environmental stress and pollution from anthropogenic sources, aiding restoration efforts in adequately diagnosing issues and improving the health and cycling patterns of restored reefs.

Thus, restored sites host a mixture of host-associated, sediment-associated, and marine free-living taxa (Fig.4), consistent with the multi-habitat coupling typical of oyster reefs (Hernández-Zárate & Olmos-Soto, 2006). This pattern, together with evidence of active nutrient cycling and detectable AMR determinants that may constrain *Vibrio* and other opportunists, suggests that microbiome-aware restoration can deliver co-benefits for reef function, environmental protection, and human health.

## 5. IMPLICATIONS FOR MONITORING AND MANAGEMENT

Building on these findings, we recommend a microbiome-integrated monitoring framework (Table 2) through operationalizing three principles: (i) stratify by matrix (oyster tissue, biofilm, sediment, water), because matrix, not geography, drives community structure; (ii) prioritize biofilm resistome tracking, as biofilms concentrate AMR potential; and (iii) target *Vibrio* surveillance in oysters, where *Vibrio*-linked markers are most evident despite low overall abundance. This design aligns ecological performance metrics with public-health safeguards and provides clear triggers for adaptive management at restoration sites.

We propose the following steps for restoration management and design

### 1) Matrix-stratified sampling design

Monitor each restoration site by matrix rather than treating sites as homogeneous units. Minimum core set per campaign: oyster tissue (n≥5 pooled), biofilm scrapes (n≥3), surface water (n≥3), and surficial sediment (n≥3). Sample seasonally to capture wet/dry contrasts.

### 2) Biofilm-focused AMR panel

Use short-read metagenomics or targeted qPCR/dPCR to track a sentinel resistome panel in biofilms (e.g., tet(C), APH(3⍰)-Ia, acrAB/-D, mdtABC/H/F/P, mdfA, MexF/MexW). Treat biofilms as early-warning “hotspots”; flag step-ups in abundance or diversity of AMR determinants for follow-up source investigation (e.g., stormwater or upstream inputs).

### 3) Targeted *Vibrio* surveillance in oysters

Combine metagenomic detection with targeted assays for *V. parahaemolyticus*/*V. vulnificus* and selected virulence/resistance markers (e.g., tdh/trh, varG). Where feasible, pair with cultivation and plaque assays to resolve *Vibrio*–phage dynamics.

### 4) Decision rules and integration

Define matrix-specific baselines and trigger points (e.g., percentile thresholds or fold-changes from site baselines) that initiate adaptive actions: intensified sampling, source tracing, or temporary harvest advisories in collaboration with managers. Report indicators alongside traditional metrics (reef structure, recruitment) to align ecological and health outcomes.

### 5) QA/QC and comparability

Standardize field volumes, extraction protocols, and bioinformatics (including host-read filtering); include negative controls and internal standards to permit between-year and between-site comparisons.

In sum, a matrix-aware design with biofilm resistome tracking and oyster-focused *Vibrio* surveillance operationalizes microbiome science for restoration, providing managers with practical indicators that connect reef function to human-health protection.

